# Correlating the differences in the receptor binding domain of SARS-CoV-2 spike variants on their interactions with human ACE2 receptor

**DOI:** 10.1101/2022.09.30.510287

**Authors:** Gokulnath Mahalingam, Porkizhi Arjunan, Yogapriya Periyasami, Ajay Kumar Dhyani, Nivedita Devaraju, Vignesh Rajendiran, Abhisha Crystal Christopher, Ramya Devi KT, Immanuel Darasingh, Saravanabhavan Thangavel, Mohankumar Murugesan, Mahesh Moorthy, Alok Srivastava, Srujan Marepally

## Abstract

Spike protein of SARS-CoV-2 variants play critical role in the infection and transmission through its interaction with hACE2 receptor. Prior findings using molecular docking and biomolecular studies reported varied findings on the difference in the interactions among the spike variants with hACE2 receptor. Hence, it is a prerequisite to understand these interactions in a more precise manner. To this end, firstly, we performed ELISA with trimeric spike proteins of Wild (Wuhan Hu-1), Delta, C.1.2 and Omicron variants. Further, to study the interactions in a more specific manner by mimicking the natural infection, we developed hACE2 receptor expressing HEK-293T cell line and evaluated binding efficiencies of the variants and competitive binding of spike variants with D614G spike pseudotyped virus. In lines with the existing findings, we observed that Omicron had higher binding efficiency compared to Delta in both ELISA and Cellular models. Intriguingly, we found that cellular models could differentiate the subtle differences between the closely related C.1.2 and Delta in their binding to hACE2. From the analysis in receptor binding domain (RBD) revealed that a single common modification, N501Y, present in both Omicron and C.1.2 is driving the enhanced spike binding to the receptor and showed two-fold superior competitive binding than Delta. Our study using cellular model provides a precise method to evaluate the binding interactions between spike sub-lineages to hACE2 receptors and signifies the role of single common modification N501Y in RBD towards imparting superior binding efficiencies. Our approach would be instrumental in understanding the disease progression and developing therapeutics.

**Author Summary:** Spike proteins of evolving SARS-CoV2 variants demonstrated their signature binding to hACE2 receptor, in turn contributed to driving the infection and transmission. Prior studies to scale the binding efficiencies between the spike variant and the receptor had consensus in distinct variants, but discrepancies in the closely related ones. To this end, we compared spike variants-receptor interactions with ELISA, from cells expressing hACE2 receptor. Intriguingly, we found that cellular models could differentiate the subtle differences between the closely related C.1.2 and Delta in their binding to hACE2. More importantly, competitive binding studies in presence of pseudovirus, demonstrated that a single common modification, N501Y, present in both Omicron and C.1.2 showed two fold superior competitive binding than Delta. Collectively, our study suggests a precise approach to evaluate the binding interactions between spike sub-lineages to hACE2 receptor. This would be instrumental in understanding the disease progression and developing therapeutics.

## Introduction

The severe acute respiratory syndrome coronavirus 2 (SARS-CoV-2) has been continuously evolving into new variants by mutations, resulting into multiple waves of coronavirus disease 2019 (COVID-19) pandemic in human populations worldwide [2]. The entry mechanism of SARS-CoV-2 into host cells is primarily driven by binding of trimeric spike glycoprotein through receptor-binding motif (RBM) to cognate host receptor (angiotensin-converting enzyme 2 (ACE2)) followed by receptor mediated endocytosis [3]. Several new variants harbor single or group of mutations in altered spike proteins interaction with hACE2 which make them more infectious, transmittable, and the dominant strain during COVID-19 pandemics [4]. The first noticeable variant of SARS-CoV-2 was D614G (Pango lineage: B.1, clade 20A) with single point mutation (substitution of aspartic acid (D) with glycine (G)) in spike protein was identified beginning of 2020. The D614G showed high affinity to hACE2 by increasing open conformation of RBD (receptor-accessible state) in spike protein then D614 variant (wild) with enhanced cellular and host tropism, resulting in more infectious, transmittable, and the dominant pandemic strain that replaced D614 at mid of 2020 [5,6]. Later, several globally dominant strains have been evolved periodically with multiple mutations from D614D strains at time which replaced its predecessors and became dominant globally. The WHO has classified these variants named Alpha, Beta, Gamma, Delta, and Omicron, among others as variants of concern (VOC) [2].

The outbreaks of Delta and Omicron variants were dominant across the globe including India by breakthrough infections. Omicron is evolutionarily the most distinct VOC with a high number of mutations (36 coding mutations) in spike protein compared to other VOCs.

Due to higher transmission rate, Omicron has spread rapidly and become the dominant circulating variant globally and has replaced all other previous VOCs [7,8]. Hence, it is critical to understand the binding efficiencies of these variants to ACE2 receptor to understand potencies of infectivity and transmission rates particularly of dominant Delta and Omicron variants. Multiple computational and experimental studies were attempted to understand the impact of the mutations on binding of Delta and Omicron to ACE2 receptor. However, there were some discrepancies among these studies on the binding efficiencies of these variants. The most of computational studies predicted that RBD of Omicron showed stronger binding to hACE2 receptor compared to RBD of wild and Delta respectively [9-15]. Leyun Wu et al., reported that RBD of Omicron had a weak affinity to hACE2 receptor compared to RBD of Delta, but similar to RBD of wild affinity by molecular dynamics (MD) simulations analysis [16]. Similar findings were observed with analyzed non-competitive ELISA, in which RBD of Delta showed higher binding than RBD of Omicron [16,17]. Several attempts were made using multiple biomolecular interaction techniques such as Microscale thermophoresis (MST), bio-layer interferometry (BLI), Plasmon Resonance (SPR) to understand affinity between spike variants and hACE2 receptor. Seonghan Kim et al, demonstrated that RBD of Omicron has high affinity towards hACE2 than RBD of Delta by MST analysis. Contrasting results were found in Maren Schubert et al study [11,17]. However, BLI approach revealed that both RBDs of Omicron and Delta bind with similar affinities towards hACE2 [18]. The inconsistency in the binding affinities were also observed with SPR analysis as well [19-23]. Hence, a comprehensive strategy including comparing multiple techniques is a prerequisite for precise understanding of spike variant-hACE2 interactions.

To this end, we employed three different approaches including 1). To mimic the natural interaction, we used trimeric form of spike variants and titrated them with soluble hACE2 (using ELISA) and hACE2 expressed on cells surface using two different approaches, 2.) In-vitro flow cytometry and 3). Spike pseudovirus competitive assays.

## Materials & Methods

### Cloning of hACE gene into lentiviral transfer plasmid

The hACE2 ORF sequence amplified from hACE2 vector (Addgene, 1786) by high fidelity Q5 polymerase PCR using primer set given table-1 and cloned purified hACE2 gene fragment into NheI and BamHI digested fragment of pLenti backbone (Addgene, 112675) by Gibbson assembly. The hACE2 clones (pLenti-hACE-P2A-Puro) were confirmed by restriction digestion with NheI and BamHI enzymes.

### ELISA for hACE2 and spike variants binding

We used trimeric spike variants protein (ACROBiosystems) such as Delta (B.1.167.2), C.1.2, Omicron (B.1.1.529) were expressed in human 293 cells (HEK293) and trimeric conformation of purified proteins confirmed by SEC-MALS (as per company CoA). The hACE2 protein (SinoBiological,10108-H08H) was coated on high binding 96 Well Strip Plate (Biomat, MT01F4-HB8) at 0.1 μg per well concentration in phosphate-buffered saline (PBS, pH-7.4) overnight at 4°C. Later, plate was washed 3x with washing buffer (PBS-T, 0.05% Tween 20 in PBS) and blocked with 3% BSA in PBS-T for 2 hours at RT. After 3x washing, incubated with 100 μl of biotinylated trimeric spike protein of wild, Delta, C.1.2, Omicron variants at different concentrations (5-fold dilution in FC buffer from 5 to 0.039 μg/ml) in diluent buffer (1% BSA in PBS-T) to hACE2 coated wells for 1 hour at 37°C. After removing unbound with 5X washing, all wells were incubated with 100 μl of Streptavidin-HRP reagent diluted 1/5000 in 1% BSA in PBS-T for 1 hour at 37°C. After 5X washing, 3,3′,5,5′-Tetramethylbenzidine (TMB) substrate (Bioponda Diagnostics, TMB-S-005) was added and stopped after 5 minutes with stop solution (Bioponda Diagnostics, STP-001). Each well was read for optical density (OD) at 450 nm in i3x plate reader (Molecular Devices). OD of samples was subtracted from OD of blank (Diluent buffer only). The percentage of maximum spike binding was calculated by the following formula: [OD of concentrations /OD of highest concentration of spike variants)] × 100%. The Half maximal effective concentration (EC50) of each variant were calculated by agonist vs. normalized response --Variable slope model.

### Generation of stably expressing hACE2 HEK-293T (293T-hACE2) Cells

The hACE-P2A-Puro gene encoding VSV-G lentiviral particles were produced by second generation lentiviral plasmids (Addgene) and transfected both pLenti-hACE-P2A-Puro, psPAX2 and pMD2.G plasmids at 2:1:1 ratio into HEK-293T cells at 75-80 % cells confluency using LF-3000. After 60 h post transfection, the lentiviral particles were purified using lentivirus concentrator in supernatant and reconstituted in 40 μl DMEM media and store at -80°C. Later, transduced the purified lentiviral particles to 0.5 million HEK-293T cells in D10 media supplemented with 5 mM HEPES buffer and polybrene at 6 μg/ml concentration in a six-well plate. After 48 hours post transduction, hACE2 stable clones were established for 5 passages using puromycin at 2 μg/ml concentration in the D10 medium. The expression of hACE2 in selected clones was confirmed by qPCR, Western blot and RBD surface staining followed by flow cytometry analysis and confocal microscopy.

### qRT-PCR analysis

We used the protocol for qRT-PCR as described [24]. Total RNA was isolated from one million cells using RNA iso-plus reagent as per manufacture protocol (takara) from HEK-293T and 293T-hACE2 cells. 500 ng of total RNAs are converted into cDNA using cDNA synthesis kit (takara). Syber green-based qPCR was performed using primer sets (Table S1) with 50 ng of cDNA from each sample in the QuanStudio-6 qPCR system (Applied Biosystems) with thermocycle condition: Stage1: 95°C for 30 sec, Stage:2 followed by 40 cycles of 95°C for 5 seconds at 60°C for 30 seconds in Real-time PCR system. The Ct value of hACE2 normalized with Ct value of internal control (GAPDH gene expression) in each sample and fold change was calculated by the 2−ΔΔCT method.

### Western blot

The protein lysates were prepared from HEK-293T and 293T-hACE2 cells using RIPA buffer (Thermos scientific, VH310061) with protease inhibitor (MedchemExpress, HY-K0010) and estimated protein concentration of each sample by the Bradford assay. 20 μg of protein lysates were resolved in 10% SDS-PAGE gel and transferred onto a PVDF membrane (WH3135834). After blocking with 5% nonfat dry milk in TBST, blot was incubated with ACE2 Antibody (Thermo Scientific, MA5-32307, and dilution at 1 in 2000 in blocking buffer) and followed by Goat Anti-Rabbit IgG H&L (HRP) (Invitrogen, ab205718, dilution-1 in 10000 in blocking buffer). The blot was visualized for hACE2 band using chemiluminescence substrate in Chem-Doc imaging system (Bio-Rad). For internal control, blot was further stripped and probed for GAPDH (Biorad, MCA4740, dilution at 1 in 2000 in blocking buffer) with anti-mouse IgG antibody (H+L), Peroxidase (VectorLab, PI-2000, dilution at 1 in 10000 in blocking buffer), and visualization of bands in the blot was performed.

### Flow cytometry

2.5×10^5^ HEK-293T and 293T-hACE2 cells resuspend in 100μl of FC buffer (2% heat inactivated FBS and 0.05% NaN3 in PBS) contain 1μg /ml biotinylated RBD ligand (SinoBiological, 40592-V08H-B) for 30 minutes at 4ºC in FACS tube. The cells were washed twice with 1 ml of FC buffer and centrifuge at 1000 rpm for 5 minutes. The supernatant was discarded and then incubated with 100μl of Streptavidin-PE-CF549 conjugate 1/100 dilution in FC buffer for 30 minutes. After 2X washing, the cells were acquired using BD Celesta and mean fluorescent intensity (MFI) of each sample was analysed by FlowJo software.

### Fluorescence confocal microscopy

2×10^5^ 293T-hACE2 cells were fixed on poly-lysine coated confocal dish (35mm, IBDI) using 300μl cold 4% Paraformaldehyde for 15 minutes. After 3X washing of fixed cells with 2 ml of cold wash buffer (4 % BSA in PBS), incubated with 300μl of 1μg /ml of the biotinylated RBD ligand for 45 minutes at 4ºC and followed by staining with streptavidin-PE-CF549 conjugate at 1/100 dilution for 45 minutes at 4ºC and washed three times with wash buffer. The cells were stained with 300μl of 100 dilutions of DAPI (1mg/ml) for 15 minutes at room temperature and washed for one time followed by addition of two drops of anti-fade (VectorLab, H-1000-10) and observed under confocal microscope.

### *In vitro* flow cytometry binding assay

1×10^5^ 293T-hACE2 cells in 100 μl FC buffer were incubated with different concentration (2-fold dilution in FC buffer from 2.5 to 0.039 μg/ml) of biotinylated trimeric spike protein of Wild, Delta, C.1.2, Omicron variants for 30 minutes at 4ºC. The cells were washed twice with FC buffer by centrifugation (1000 rpm/5minutes) and incubated with streptavidin−PE conjugate (BD bioscience, 349023) for 30 minutes at 4ºC. Cells were washed with PBS, centrifuged, and reconstituted in 100 μl of FC buffer. The stained cells were acquired for PE positive cells in flow cytometry BD Celesta. Level of percentage and MFI of PE positive cells were quantified by gated from control cells (cells + streptavidin−PE conjugate). The percentage of maximum spike binding for each variant was calculated by MFI using the following formula: [MFI of each concentration /MFI of highest concentration of spike variants)] × 100%. The Half maximal effective concentration (EC_50_) of each variant were calculated by agonist vs. normalized response --Variable slope model.

### Production of G614 spike pseudotyped Lentiviruses

G614 spike pseudotyped Lentiviral particles were produced as per protocol described in [25]. Briefly, HEK-293T cells at 50–70% confluent in 100 mm plate was transfected with BEI lentiviral plasmids at concentration of 5 μg plentiviral backbone (Luciferase-IRES-ZsGreen), 1.1 μg pHMD-Hgpm2, 1.1 μg plasmid pRC-CMV-Re v1b, 1.1 μg pHDM-tat1b and 1.7 μg pG614 spike protein in 10 ml of D10 growth media (10% FBS in DMEM media) using Lipo-DOH transfection reagents. At 18–24 h post-transfection, media was replenished with 10 ml fresh D10 media and incubated at 37°C for 60 h. Later, G614 spike pseudoviruses were harvested by collecting the supernatant from plate, centrifuged, and filtered by using a 0.45 μm PVDF low protein-binding filter, and stored at −80°C for further analysis.

### In-vitro spike pseudovirus competitive assay

G614 spike pseudovirus competitive assay was performed with modification of G614 spike pseudovirus neutralization assay as described [25]. Briefly, the 293T-hACE2 cells were seeded (1.25 × 10^4^ per well) in 0.1 mg/ml poly-l-lysine-coated 96-well plate. On next day, cells were incubated 50 μL serially diluted (2-fold dilution in FC buffer from 20 to 0.039 μg/ml) of biotinylated trimeric spike protein of Wild, Delta, C.1.2, Omicron variants in D10 media and 50 μL of G614 spike-pseudoviral supernatant (∼2-4×10^6^ RLU per mL of D10 media) in respective wells at 37°C in CO2 incubator. At 60-72 h post transduction, infectivity of G614 spike-pseudovirus was visualized by expression of ZsGreen level under fluorescence microscopy (Leica), and quantified infectivity by luciferase activity using Steady-Glo® luciferase reagent (Promega, E2510). From Relative Luciferase Unit (RLU) values, the percentage of maximum G614 spike pseudovirus infectivity for each variant at different concentration was calculated by the following formula: [RLU of each concentration /RLU of highest concentration of spike variants)] × 100%. Half-maximal inhibitory concentration (IC_50_) of each variant were calculated using [Inhibitor] vs. response --Variable slope (four parameters).

### ELISA for anti-RBD antibody and spike variants interaction

The trimeric spike protein of Wuhan Hu-1, Delta, C.1.2, Omicron variants were coated on high binding 96 Well Strip Plate (Biomat, MT01F4-HB8) at 0.1 μg per well using phosphate-buffered saline (PBS, pH-7.4) overnight at 4°C. Blocked with 3% BSA in PBS-T for 2 hours at room temperature and incubated with 100 μl of anti-RBD mab (2-fold diluted in 1% BSA in DPBS-T from 2.5 to 0.02 μg/ml concentration) to spike variants coated wells for 1 hour at 37°C. All wells were washed and incubated 100 μl of goat anti-rabbit IgG specific−HRP antibody at dilution of 1 in 5000 in blocking buffer. The level of anti-RBD bound to each spike variants were estimated by 3,3′,5,5′-Tetramethylbenzidine (TMB) substrate. The level of binding of IgGs in each sera sample to spike variants were quantified by measuring the optical density (OD) at 450 nm in i3x plate reader (Molecular Devices, San Jose, CA). OD of each sera sample for spike variants was subtracted to respective spike variants blank OD (without sera). The percentage of maximum anti-RBD mab binding was calculated by the following formula: [OD of concentrations /OD of highest concentration of spike variants)] × 100%. The Half maximal effective concentration (EC_50_) of each variant were calculated by agonist vs. normalized response --Variable slope model.

### Statistical analysis

The statistical significance and visualization of data were generated in GraphPad Prism version 8.0. Statistical significance of spike variant binding to hACE2 receptor was analyzed by unpaired t test with Welch’s correction. Equal or less than p values, of *p < 0.05, **p < 0.01, considered as significant and ns = non-significant.

## Results

Towards understanding the differences in the binding affinities of spike variants including Wild, Delta, Omicron and C.1.2 with hACE2 receptor, we employed three different methods including ELISA, in vitro flow cytometric binding and competitive binding assay. We also followed three parameters for this study (i) To mimic natural forms of spike protein on SARS-CoV-2 variants, we used trimeric form of spike variants (mutational profile of each variant given in Figure 1) rather than RBD domain to assess the binding activity towards human ACE-2 (hACE-2) receptor (ii) The differential mutational profiles of spike variants affect the molecular weight of each spike protein of variants, for homogeneity. Hence, we evaluated binding efficiencies of individual spike variants towards hACE2 receptor in molarity, (iii) Binding efficiency of individual spike variants is compared between monomeric as well as dimeric (on cell surface) form of hACE2 receptor.

**Figure 1.**
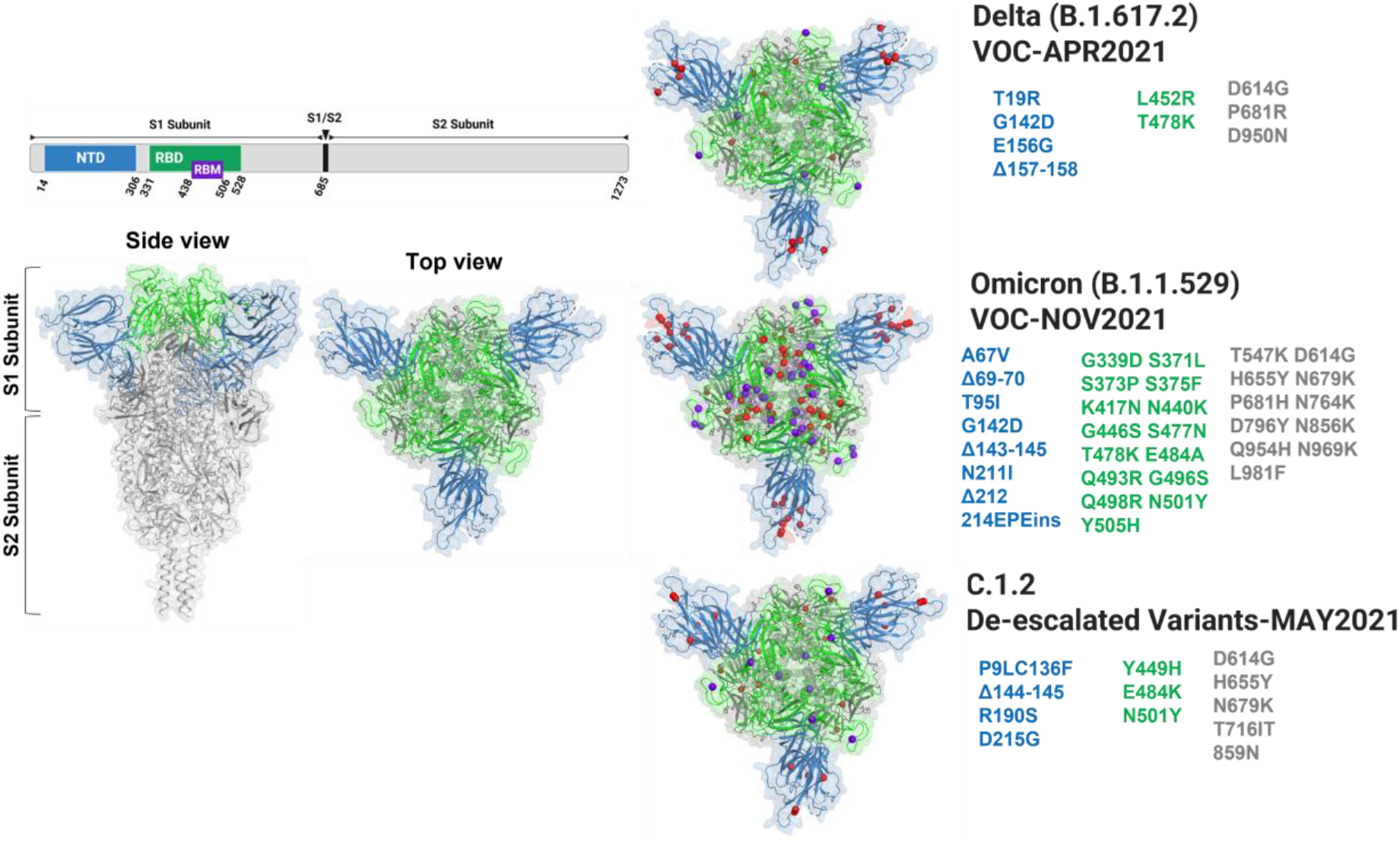
3D crystal structure of ancestral SARS-CoV-2 spike trimer (PDB ID 6XR8) [1] was generated, highlighting the mutational landscape by purple balls in RBM domain, red balls in NTD (blue), RBD (green) and other (gray) regions of Delta, Omicron and C.1.2 spike trimers using PyMOL software.

Firstly, we evaluated the binding efficiency of trimeric spike variants towards soluble hACE2 receptor using ELISA based assay (Figure 2A). We calculated the binding efficiency of 50 (EC_50_) by concentration-response curves for binding of SARS-CoV-2 spike variants to hACE2 using % of maximum ligand binding values. All spike variants were efficiently binding to hACE2 protein compared to wild spike. Among the spike variants, Omicron showed 6-fold stronger binding towards hACE2 (EC_50_-0.38 nM), than Delta (4.8-fold, EC_50_-0.48 nM) and C.1.2 (3.7-fold, EC_50_-0.61 nM) when compared to Wild spike (Figure 2B). From these findings, Omicron showed stronger binding towards hACE2 receptor when compared to Delta and C.1.2 variants.

**Figure 2.**
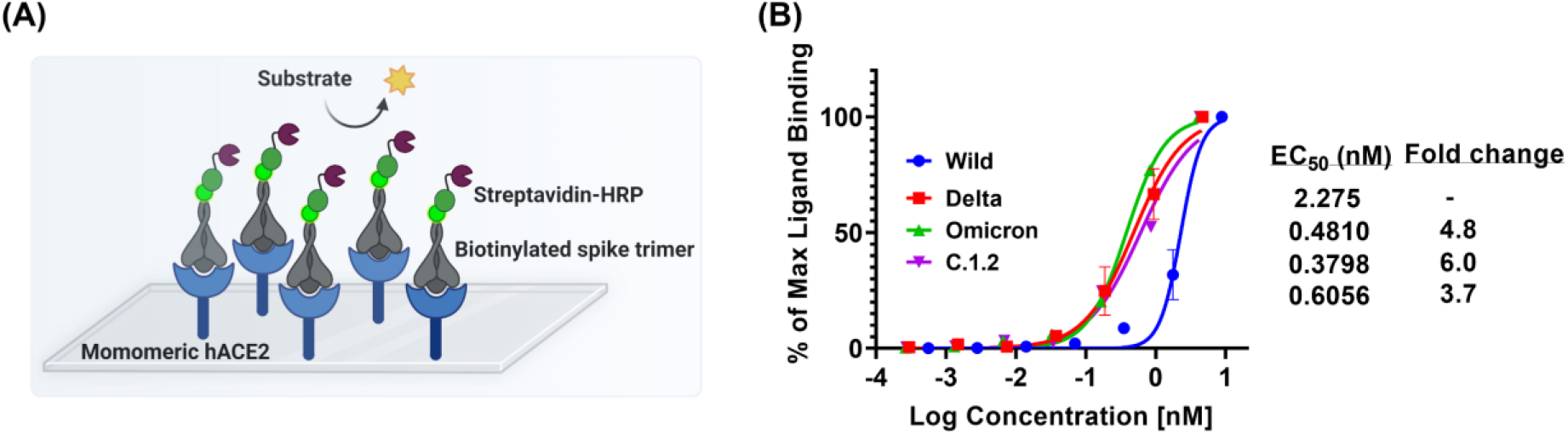
Concentration−response curves of trimeric SARS-CoV2 spike protein variants, binding to hACE2 receptor protein. Schematic representation of ELISA assay (A); The binding curves and EC50’s of each spike variants was quantified using ELISA by titrating of biotinylated spike variants to soluble hACE2 coated on the plate and the amount of bounded spike variants were estimated using streptavidin−HRP conjugate (B). (Mean ± SEM, N=2).

To closely mimic the natural infection, hACE2 receptor was expressed on the surface of HEK293 cells and evaluated the binding efficiency of these variants using in-vitro flow cytometry binding assay (Figure 2A). Firstly, hACE2 gene was cloned into lenti-backbone and produced lentiviral particles encoded hACE2 receptor (Figure S1). These particles were transduced into HEK293T cells for generation of stable hACE2 overexpressing cells (293T-hACE2). The expression of hACE2 on 293T-hACE2 cells was confirmed by qPCR, showed overexpression of hACE2 mRNA (∼7000-fold) and western blot, respectively (Figure 3A&B). Further, we confirmed surface expression of hACE2 receptor on the cells by flow cytometry and fluorescent confocal microscopy (Figure 3D-F).

**Figure 3:**
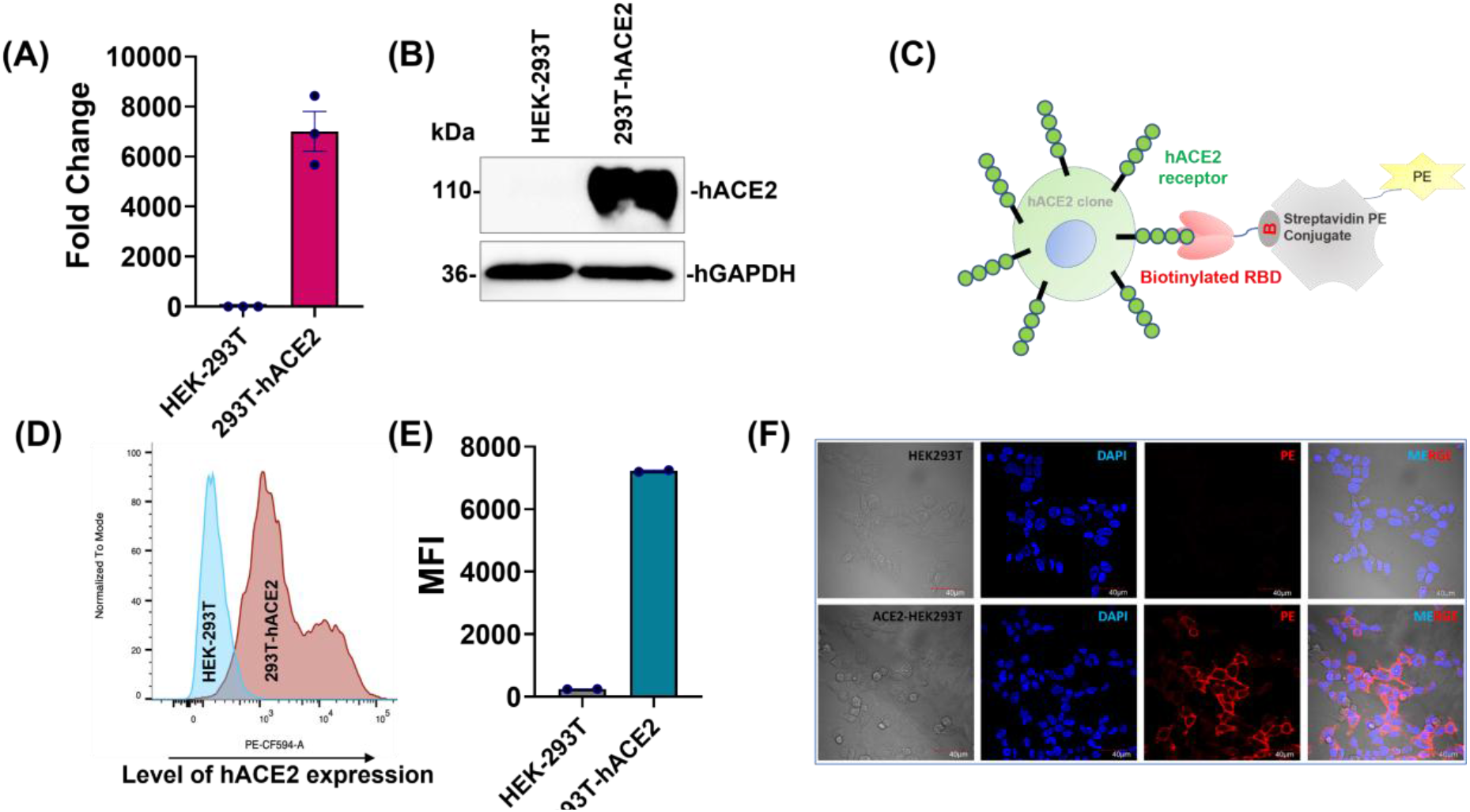
Generation and characterisation of hACE2 receptor stably expressing HEK-293T cell line. The VSV-G lentiviral particles were produced with pLenti-hACE2-P2A-PuroR plasmid and transduced into HEK-239T cells. The stable expressing hACE2 HEK-239T (293T-hACE2) cell line was selected with puromycin. The expression level of hACE2 in 293T-hACE2 cells were analysed by Qpcr (A) and western blotting techniques (B). The functional characterization of 293T-hACE2 stable cells were done by RBD-Biotin surface staining. Graphical representation of RBD-biotin surface staining principle (C). After RBD-Biotin surface staining, the ancestral SARS-CoV-2 RBD binding hACE2 level on surface of 293T-hACE2 cells were quantified by flow cytometry (D-E) (MFI-mean florescent intensity) and RBD-hACE2 interaction was visualised by confocal microscopy (F).

Using these 293T-hACE2 cells, we performed *in vitro* flow cytometry binding assay at different concentrations of trimeric spike variants (Figure 4A). Although the percentage of spike bound cells was similar among Wild, Omicron, Delta, and C.1.2 spike-stained cells (Figure 4B & S2), the level of spike variants binding (MFI) on 293T-hACE2 cells was higher in Omicron, Delta, and C.1.2 spike-stained cells compared to wild spike-stained cells (Figure 4C & S3). The fold change of spike binding was calculated by normalizing to wild spike. We found that around 2.5 to 1.3-fold increase their binding on stained cells at higher concentration of other spike proteins compared to Wild (Figure 4D). Further, we analyzed the EC_50_ by concentration-response curves using % of maximum ligand binding values (MFI) between Delta, Omicron and C.1.2 spike proteins. When compared to Delta (EC_50_-1.2 nM) and C.1.2 spike (EC_50_-1.1 nM), Omicron spike showed stronger binding to hACE2 on surface of the cells (EC_50_-0.78 nM) (Figure 4E). It indicated that Omicron showed stronger binding to dimeric hACE2 receptor cells, which is consistent with above data. Moreover, this data revealed that binding efficiencies of both Delta and C.1.2 were similar to hACE2 receptor in its native form on the cells.

**Figure 4.**
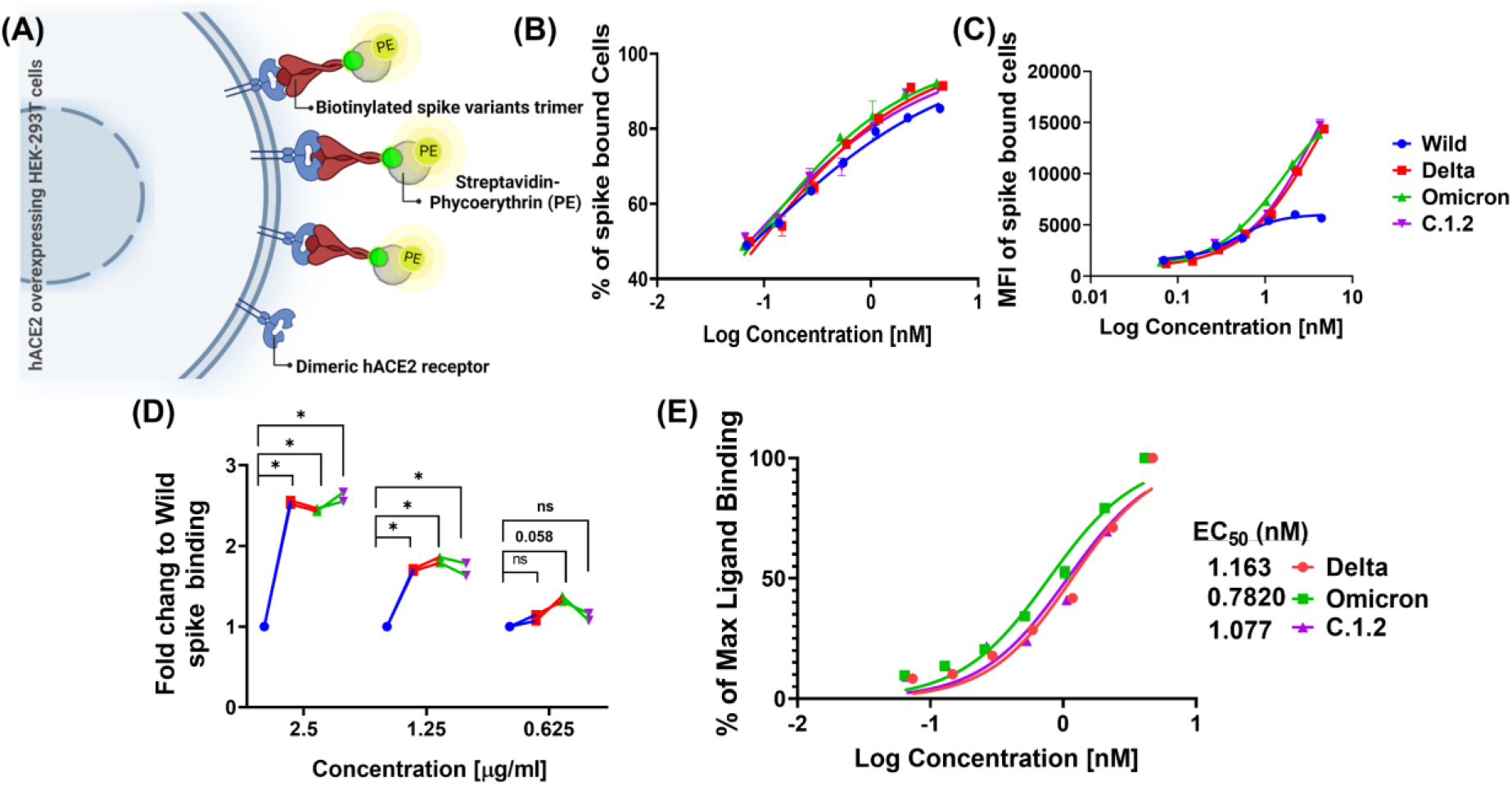
Potency of spike variants affinity to dimeric hACE2 receptor present on surface of the cells. Schematic representation of *in vitro* flow cytometry binding assay to estimate binding affinity of spike variants (A). The 293T-hACE2 cells (hACE2 receptor overexpressing HEK-293T cells) were surface stained with biotinylated trimeric spike variants at different concentration. Percentage of spike variants bounded cells (B) and level of binding (MFI) (C) on spike bounded cells quantified using streptavidin−PE conjugate in flow cytometry. The fold change of binding (MFI) for each spike variants related to wild calculated at indicated concentration (D). Percentage of maximum spike binding was calculated from MFI, plotted against concentration and a non-linear curve fitting was used to estimate the EC_50_ of spike variants (E).

To confirm further, we also analyzed binding efficiencies of spike variants towards hACE2 receptor in the presence of spike G614 mutant pseudovirus at different concentrations of spike variants (competing with G614 spike pseudovirus) in 293T-hACE2 cells (Figure 5A) [4]. The pseudovirus has zsGreen encoding gene, upon the infection, zsGreen expression is observed. The inhibition of G614 spike pseudovirus (zsGreen expression) can be analyzed in fluorescent microscopy. We observed that Omicron, Delta, and C.1.2 spike protein inhibited the G614 spike pseudovirus infectivity more efficiently than wild spike protein in 293T-hACE2 cells (Figure 5B). The strength of binding efficiency of each spike variants was calculated by inhibitory concentration (IC) value. The IC value is inversely correlated with binding affinity of spike variants (i.e., lower IC_50_ or IC_90_ value > higher affinity). We also used luciferase gene encoding G614 spike pseudovirus in a competitive binding assay for highly sensitive detection. Luciferase expression is directly proportional to the pseudovirus infection. Consistent with above two methods, spike protein of Omicron showed higher affinity (IC_50_-1.1 nM) towards hACE2 receptor on 293T-hACE2 cells compared to Delta (IC_50_-1.4 nM), C.1.2 (IC_50_-1.3 nM) and wild variant (IC_50_-7 nM) (Figure C-F). The receptor affinity was increased to spike of Omicron (6.2-fold) Delta (5.1-fold) and C.1.2 (5.4-fold) when compared to wild spike (Figure 5G). Since mild differences in the affinities of Delta and C.1.2 spike to hACE2 dimer were observed, we used IC_90_ (inhibit 90 % of G614 spike pseudovirus infectivity at concentration of spike variants) to understand in a more precise manner. Interestingly, we found that the binding affinities of Omicron (3-fold) and C.1.2 (2.5-fold) to hACE2 dimer were higher compared to Delta (Figure 5G). Overall, the receptor binding studies in cellular model revealed an increasing affinity of spike proteins to hACE2 receptor from Omicron>C1.2>Delta>Wild variant.

**Figure 5.**
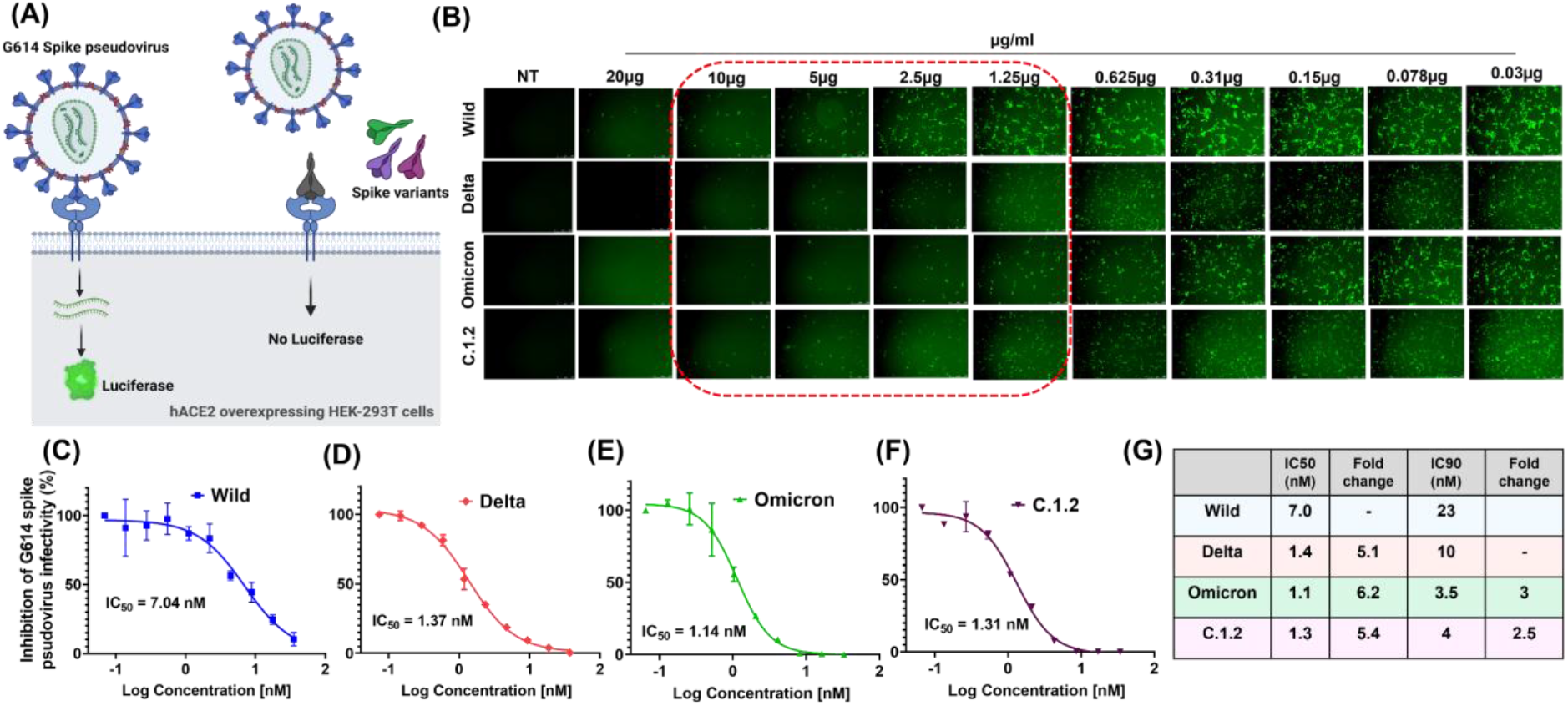
Probing spike variants affinity to dimeric hACE2 receptor expressing on surface of the cells by competing G614-spike pseudovirus infectivity with spike variants. Schematic representation of *in vitro* competitive pseudovirus assay (A). The 293T-hACE2 cells was incubated with G614-spike pseudovirus (expressing luciferase and ZsGreen protein) and different concentration of spike variants. After 60-72 hours, infectivity of G614-spike pseudovirus was visualized by expression of ZsGreen protein in fluorescent microscopy at different spike variant treated conditions (B). The percentage of G614-spike pseudovirus infectivity was quantified by luciferase activity and inhibitory concentration 50 (IC_50_) of wild (C) Delta (D) Omicron (E) C.1.2 (F) spike proteins were quantified using non-linear curve fitting models. (G) Table of IC50 and IC90 values of each spike variants

Our findings in both ELISA based assays and cellular models collectively demonstrated that mutational profile on RBD domain of spike variants increased their binding towards hACE2 receptor. Combination of spike, affinity to hACE2 receptor and immune evasion are involved in establishing a new variant in human populations. Hence, we also analyzed whether the mutations on RBD domain of spike variants affect its antibody binding. To this end, we screened anti-RBD monoclonal antibody (raised against RBD domain of Wild spike) binding to spike variants by ELISA (Figure 6A). We observed that anti-RBD mab showed higher affinity to spike protein of wild (EC_50_-0.13 μg/ml), but affinity was significantly reduced to spike proteins of Delta (EC_50_-0.66 μg/ml), Omicron (EC_50_-0.52 μg/ml) and C.1.2 (EC_50_-0.54 μg/ml) variants (Figure 6B). Collectively, these studies revealed that Omicron and C.1.2 had higher affinity compared to Delta and wild type.

**Figure 6.**
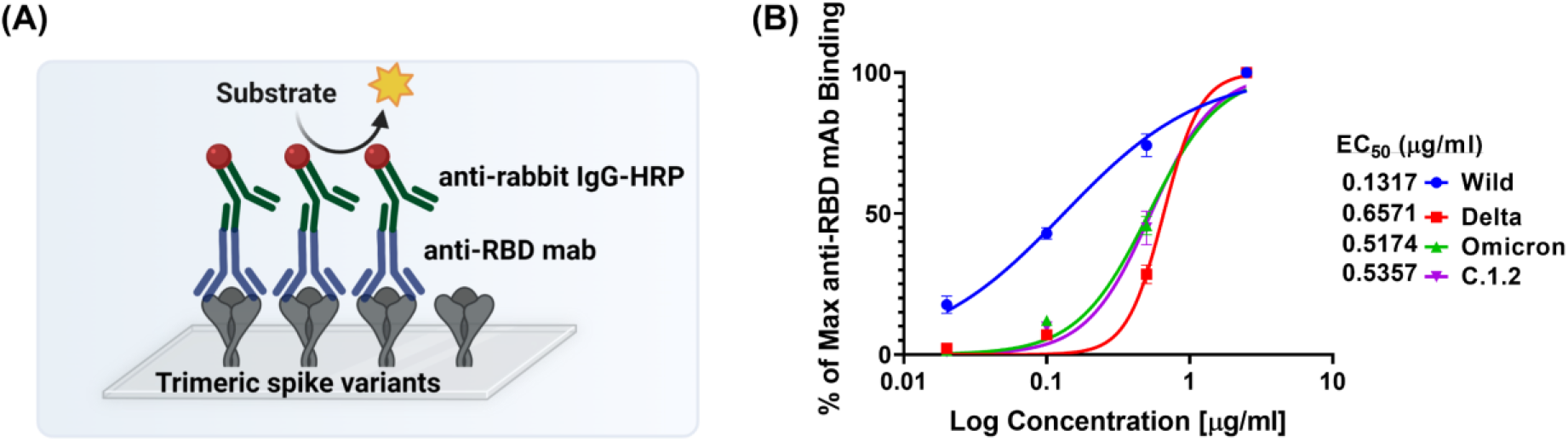
Concentration−response curves of anti-RBD antibody binding to trimeric SARS-CoV2 spike variants protein. Schematic representation of ELISA assay to quantify the binding of anti-RBD antibody against trimeric SARS-CoV2 spike variants in fluorescent microscopy (A). EC50 was obtained by titrating of anti-RBD antibody against trimeric spike protein coated on the plate and level of bounded antibodies of each spike variants were quantified using streptavidin−HRP conjugate (B) (mean ± SEM, N=3).

## Discussion

Understanding the interaction between spike protein variants and hACE-2 receptor is critical for evaluating the infectivity and transmission rates of the VOCs. Most of the studies including molecular modelling studies, biomolecular interaction studies and ELISA are confined to RBD domain, not to the stable homodimer form of the hACE2 receptor. Since, recombinantly produced soluble hACE2 dimer (1-740 aa residues) is less stable, and can be converted into monomeric form, hence it is stabilized with dimeric FC domain [26,27]. The critical challenge in understanding the RBD of spike and hACE2 interactions are limited by the techniques are being used.

Evaluating binding energies of trimeric spike to the native form of the receptor using molecular modeling simulations is limited by larger size [28]. SPR and BLI based method need chemical modifications on the biomolecules for attaching onto the surface of chip [16]. As there is an inconsistent data with these techniques from different groups, we anticipate that these chemical modifications on RBD of spike protein may have potential interference in determining their affinities to the receptor and spike trimer function different from RBD.

Considering the limitations in these techniques, we compared 3 different techniques for a precise evaluation of binding efficiencies. To this end, firstly, we employed non-competitive ELISA, coated with soluble hACE2 receptor, and evaluated the binding efficiencies of trimeric spike variants from Wild, Delta, C.1.2 and Omicron. The trimeric spike of omicron has 6-fold highest affinity toward soluble hACE2 receptor among Delta (4.8-fold), C.1.2 (3.7-fold) than wild.

Recent findings reported that homodimeric forms of hACE2 either in soluble or on cell surface enhanced the spike protein binding than monomeric forms and disruption in the dimerization significantly affected the interactions [26,27,29,30]. Hence, it is ideal to evaluate the receptor binding interactions of spike variants in a close to natural phenomenon. It is observed that there are discrepancies in the results within and among the techniques led to inconclusive experimental evidence.

To the best our knowledge, so far there are no attempts to evaluate and correlate the binding efficiencies of spike variants in cellular models. To this end, we studied the binding efficiencies of spike proteins with homodimeric hACE2 receptor, expressed on the surface of the cells using biotinylated trimeric spike variants followed by fluorescently conjugated streptavidin binding and evaluated the fluorescent intensity in flow cytometry. Consistence with ELISA using soluble hACE2 receptor data, the affinity of trimeric spike of Omicron showed higher affinity towards hACE2 receptor on the cells than Delta and C.1.2., but affinity of trimeric spike of C.1.2 variants showed slightly higher than Delta variant towards hACE2 receptor on cell surface. We further compared these interactions with a competing with a zsGreen and luciferase expressing genes encapsulated SARS-CoV2 spike G614 pseudovirus, revealed the trimeric spike of Omicron (3-fold) and C.1.2 (2.5-fold) variant was found to be higher when compared to Delta variant.

Recent findings revealed that the spike N501Y mutation containing variants, have higher affinity to hACE2 receptor [31,32]. In concurrence, we observed similar binding affinity of N501Y mutation containing variants including Omicron and C.1.2 was increased to hACE2 receptors than Delta (does not containing N501Y mutations) by cellular model. Based on our data, rapidly expanding variant (Omicron) showed higher affinity than the C.1.2., possibly because of additional mutations such as S477N and Q498R in Omicron variants, known to strengthen the affinity of hACE2 as well as cross-species ACE2 [32].

## Conclusions

This study provides a precise approach to evaluate the binding strength of spike to cognate receptors. Using two different approaches involving cellular models, we could identify the subtle differences between the closely related C.1.2 and Delta in their binding to hACE2. We further confirmed that single modification in the RBD domain N501Y enhances the binding efficiency to the receptor. We anticipate that this approach could be considered for evaluating the binding efficiencies of emerging variants to understand the virus-host interactions, disease progression and instrumental in developing therapeutics.

## Acknowledgements

MS thanks Department of Biotechnology, India, for the financial support through grants, BT/PR25841/GET/119/162/2017; BT/PR40446/COV/140/5/2021; and Dr. Sandhya Rani for FACS and Imaging facility at CSCR-Vellore. We thank Dr. VGM Naidu, NIPER-Guwahati for his inputs in calculating binding efficiencies.

## Author contribution statement

GM, and MS contributed to the conceptualization of the article. GM, RDKT, YP, PA, AKD, ND, VR, and ACC contributed to the methodology of the article. GM, contributed to interpretation, and visualization of data. MS supervised the investigation. ID for molecular visualization of Spike variants. GM and MS contributed to manuscript preparation. GM, AS, SBT, MKM, MM and MS contributed to reviewing and editing the manuscript. All authors had access to the data. GM and MS accessed the original data and vouch for its authenticity.

